# Unified down-stream analysis of crosslinking mass spectrometry results with pyXLMS

**DOI:** 10.64898/2025.12.18.695169

**Authors:** Micha J. Birklbauer, Louise M. Buur, Sabrina Kaser, Fränze Müller, Manuel Matzinger, Karl Mechtler, Stephan Winkler, Viktoria Dorfer

## Abstract

Crosslinking mass spectrometry has become the method of choice for the identification of protein–protein interactions and for gaining insight into the structures of proteins *in vivo*. However, connecting crosslink search engine results with down-stream analysis tools, and therefore gaining biological insight from crosslink identifications, has remained a manual and cumbersome step in the analysis that often requires expert bioinformatics knowledge. Here we introduce pyXLMS, a python package and public web application which aims to simplify and streamline this intermediate step, enabling researchers even without bioinformatics knowledge to conduct in-depth crosslink analyses. In its current state pyXLMS supports input from seven different crosslink search engines, as well as the mzIdentML format of the HUPO Proteomics Standards Initiative. Down-stream analysis is facilitated by functionality that is directly available within pyXLMS such as aggregation, validation, annotation, filtering, and visualization. In addition, the data can easily be exported to more than ten supported down-stream analysis tools and formats. We demonstrate the applicability and benefits of pyXLMS by re-analyzing a publicly available crosslink dataset with a variety of different search engines and show how the same data analysis workflow can be applied using pyXLMS. pyXLMs is available via https://github.com/hgb-bin-proteomics/pyXLMS.

## 1 Introduction

Crosslinking mass spectrometry (XLMS) has matured into a powerful tool for structural, molecular, and systems biology [1] and has become the method of choice for gaining insight into the structures of proteins *in vivo* [2, 3] and the identification of protein–protein interactions [4, 5] potentially up to system-wide scale. Advancements and optimizations in crosslinking chemistry [6–8], experimental procedures [9–11], instrumentation [12], and the release of numerous robust crosslink search engines [13– 15] have made crosslink identification increasinly more appealing even for complex samples and proteome-wide searches. Plenty of comprehensive reviews have been published in recent years highlighting successful XLMS applications, potential pitfalls and drawbacks [16–22]. However, while XLMS has seen substantial progress and become more and more of a routine problem with established workflows, connecting crosslink search engine results with down-stream analysis tools such as AlphaLink2 [23], ProXL [24, 25], xiVIEW [26], or visualization tools like PyMOL [27] and ChimeraX [28–30] has remained a manual and cumbersome step in the analysis. It often requires extensive data re-shaping or transformation that is only possible for scientists with a strong bioinformatics background and yet this step is crucial for gaining biological insight from the identified crosslinks.

Even though the mzIdentML format [31] developed by the HUPO Proteomics Standards Initiative has introduced support for crosslinks already in version 1.2 [32] in 2017 and further extended the format specifically for the reporting of crosslink results recently in version 1.3 [33], the format has not seen wide adoption in the field of XLMS with only a handful of crosslink search engines and down-stream analysis tools supporting the format. Most crosslink search engines report results at different aggregation levels, for example at the crosslink-spectrum-match (CSM), crosslink/residue pair, or protein-protein interaction level, in some kind of tabular format. This is similar for down-stream analysis tools that usually require one of the aggregation levels, in most cases either CSM or residue pair level, also mostly in tabular formats for further processing. Although some down-stream analysis tools also implement their own parsers for reports of certain crosslink search engines, researchers are mostly left to their own accords when it comes to connecting crosslink identifications from search engines to down-stream analysis tools. A process that is not only time-consuming and requires bioinformatic knowlege, but also error prone.

In the following we present pyXLMS, a python package and web application with graphical user interface that aims to simplify and streamline this intermediate step, enabling scientists even without bioinformatics knowledge to conduct in-depth crosslink analyses and shifting the focus from data transformation to data interpretation. In its current state pyXLMS supports input from seven different crosslink search engines, namely MaxLynx [15] (part of MaxQuant [34]), MeroX [35, 36], MS Annika [13, 37, 38], pLink 2 and pLink 3 [39], Scout [14], xiSearch [40] and the validation tool xiFDR [41], XlinkX [42], as well as the mzIdentML format [31–33] of the HUPO Proteomics Standards Initiative. Moreover, pyXLMS also supports a well-documented and human-readable custom tabular format that allows for parsing of any generic table. Down-stream analysis is facilitated by functionality that is directly available within pyXLMS such as aggregation, validation, annotation, filtering, and visualization of CSMs and crosslinks. In addition, pyXLMS enables easy export to the required data format of various available down-stream analysis tools such as AlphaLink2 [23], ProXL [24, 25], xiNET [43], xiVIEW [26], xiFDR [41], XlinkDB [44–47], xlms-tools [48], PyMOL [27](via PyXlinkViewer [49]), ChimeraX [28–30](via XMAS [50]), or IMP-X-FDR [51].

We demonstrate the applicability and benefits of pyXLMS by re-analyzing a publicly available crosslink dataset and show how the same data analysis workflow can be applied across several different crosslink search engines using pyXLMS, eliminating the need for manual adoption to the various data formats. pyXLMS is publicly available under a permissive open-source license via https://github.com/hgb-bin-proteomics/pyXLMS and welcoming contributions by the community with the aim of supporting as many crosslink search engines and down-stream analysis tools as possible. We are also hosting a publicly available and free of charge web server that includes most functionality at https://hgb-bin-proteomics.github.io/pyXLMS-app.

## 2 Results

### 2.1 pyXLMS - a python package to streamline crosslinking data analysis

Down-stream analysis of crosslinking results and therefore gaining meaningful biological insights, requires the connection of crosslink identifications from crosslink search engines with down-stream analysis tools. This process often involves extensive reshaping, filtering and transformation of the originally reported data of the crosslink search engine, and an in-depth understanding of the chosen down-stream analysis tools is needed to successfully make this connection. We here present pyXLMS, a python package that we implemented to eliminate this intermediate process of connecting crosslink search engines with down-stream analysis tools. An overview of the package is given in Figure 1.

**Fig. 1.**
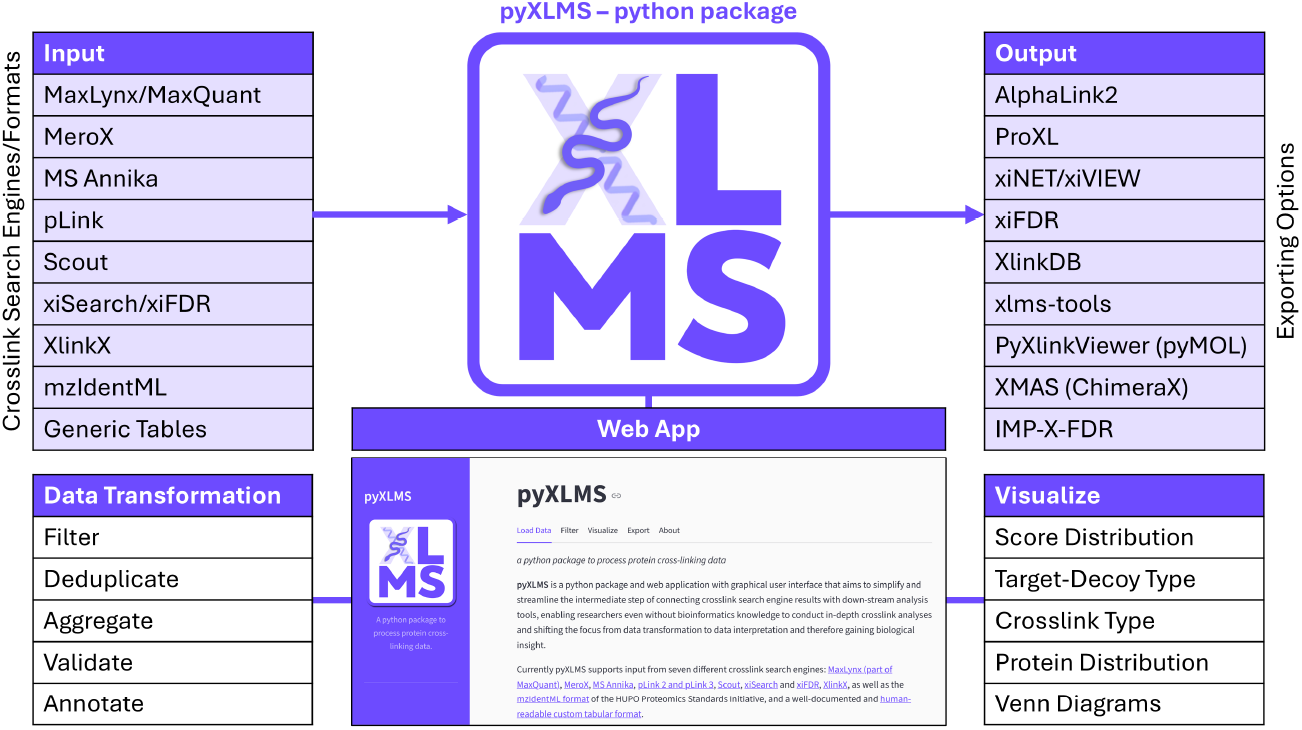
Overview of the pyXLMS python package. pyXLMS currently supports input from seven different crosslink search engines as well as the mzIdentML format. Crosslinking data can then be transformed via different filtering, annotation, and validation options or visualized with one of readily available plotting functionalities. Most importantly, the crosslinking data can be exported to the required data format of more than ten different down-stream analysis tools, eliminating the need for manual data re-shaping and transformation. pyXLMS is available as a python package and as a web application with graphical user interface.

pyXLMS handles reading of crosslink search engine results at both crosslink-spectrum-match (CSM) level and crosslink/residue pair (XL) level and currently supports seven different crosslink search engines as well as the mzIdentML format [31– 33] as input. Furthermore, pyXLMS is generally able to read any crosslink output file even of non-supported crosslink search engines with minimal additional user input. A full list of all supported crosslink search engines and formats is given in Table 1.

**Table 1.**
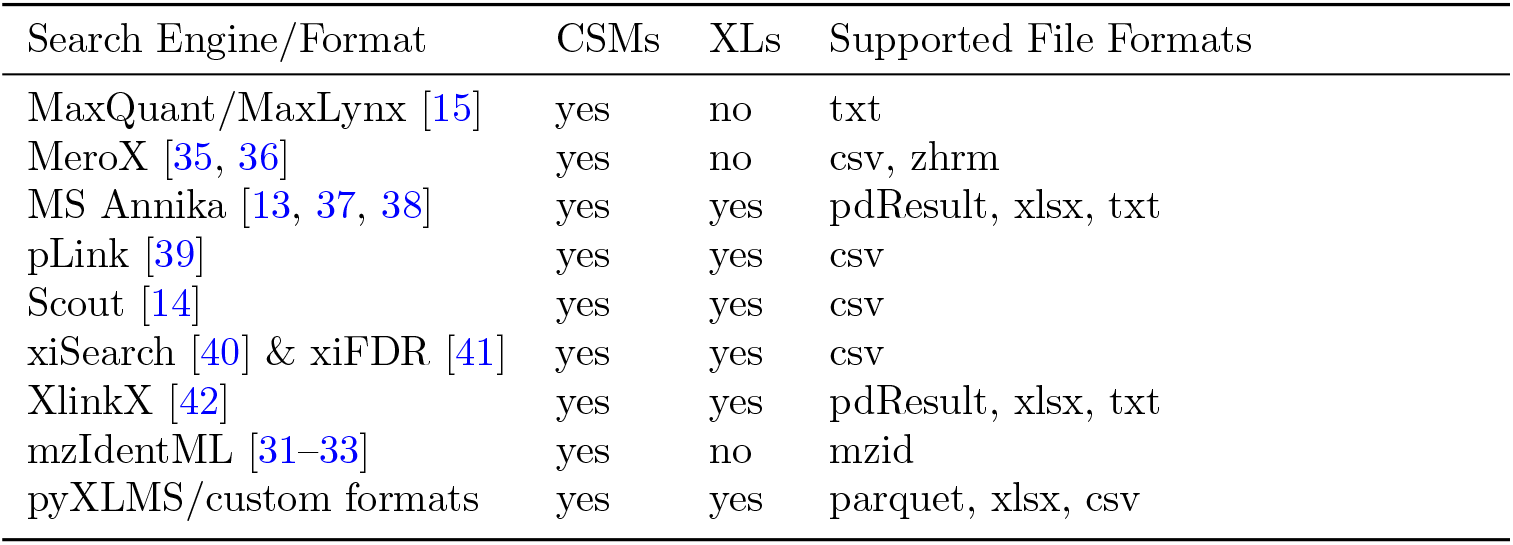
Supported crosslink search engines and formats in pyXLMS including indicators of whether the specific search engines/formats support crosslink-spectrum-matches (CSMs) and/or crosslinks/residue pairs (XLs).

Subsequently, pyXLMS allows direct inspection of the crosslinking data via a summary function that calculates common statistics, as well as several pre-defined visualization options that display score distribution, number of target and decoy matches, intra and inter matches, and the most frequently crosslinked proteins and peptides (shown in Figure 4). Data can then be filtered by target-decoy type, crosslink type, or proteins of interest. One of the most important features in pyXLMS is the ability to re-annotate protein crosslink positions based on the given XLs. Many crosslink search engines only report a single matching protein per peptide, even for ambiguous peptides that are shared among multiple proteins. This inherently leads to an incomplete list of crosslinked protein positions, which is crucial for calculating protein-protein interaction networks and understanding biological processes. The re-annotation functionality in pyXLMS matches all crosslinked peptides to the protein sequences of a given FASTA file and calculates all possible matches and protein crosslink positions, resolving this issue. Moreover, pyXLMS is able to deduplicate and filter CSMs and XLs to only contain unique matches, something that is desired for almost all down-stream analyis methods. Uniqueness for CSMs is determined by associated spectrum file and scan number while uniqueness of XLs is determined either by peptide sequence and peptide crosslink position, or by protein crosslink position, depending on user choice. pyXLMS also supports aggregation of CSMs to XLs as well as validation via false-discovery-rate (FDR) estimation. However, validation is limited to results from crosslink search engines that report both target and decoy matches, as well as a single score for FDR estimation. This is currently only the case for MaxLynx, MS Annika, and xiSearch.

pyXLMS also allows to easily create intersections of CSMs and XLs, either from different replicates or different crosslink search engines, which has been shown to increase specificity and lower the number of false positive identifications [51]. Visualization of intersections and generally overlaps of different sets of CSMs and XLs is also possible via Venn diagram plotting that is available in pyXLMS.

Last but not least, the most important feature in pyXLMS is the ability to export results to the formats needed by popular crosslink down-stream analysis tools. pyXLMS currently already supports more than ten different down-stream analysis tools and formats and handles all data re-shaping and transformation steps that are necessary. pyXLMS also implements the ProForma notation of the HUPO Proteomics Standards Initiative [52] and can convert CSMs and XLs to ProForma. A complete list of supported down-stream analysis tools is given in Table 2.

**Table 2.**
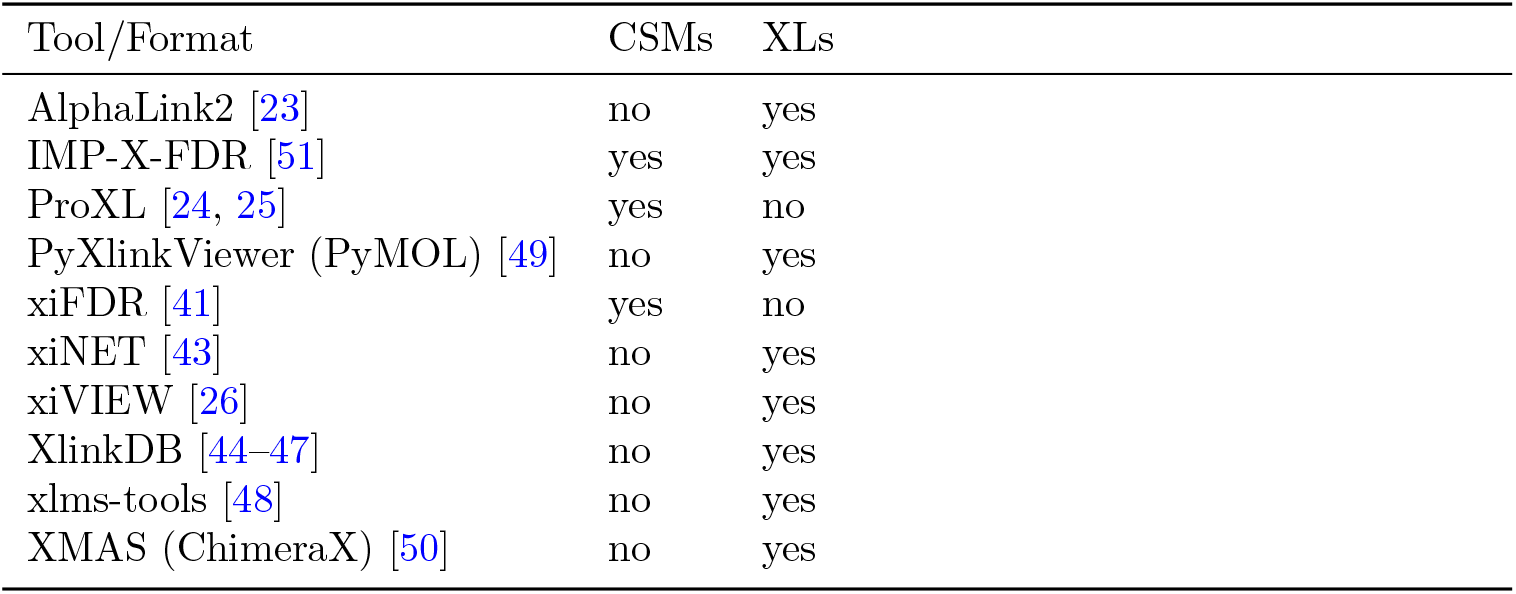
Supported export formats and down-stream analysis tools in pyXLMS including indicators whether crosslink-spectrum-matches (CSMs) and/or crosslinks/residue pairs (XLs) can be exported.

An in-depth description of the implementation of the python package is given in Section 4.1.

### 2.2 pyXLMS simplifies crosslink analysis independently of the used crosslink search engine

In order to demonstrate the applicability of pyXLMS we have re-analyzed a previously published crosslink dataset with three different crosslink search engines and show how pyXLMS unifies the down-stream analysis workflow with no need for adoption to the search engine specific result formats.

We retrieved data from PRIDE [53] where purified recombinant *S. pyogenes* Cas9 fused with a Halo-tag was crosslinked with DSBSO [54] in three replicates [11]. Subsequently, we searched the data with analogous settings using the crosslink search engines pLink [39], Scout [14], and XlinkX [42]. With pyXLMS we were able to directly load the result files from all searches, get an overview of the identified crosslinks as shown in Figure 2, and immediately continue with down-stream analysis via the export to xiVIEW [26], XMAS [50], and AlphaLink2 [23] as depicted in Figure 3.

**Fig. 2.**
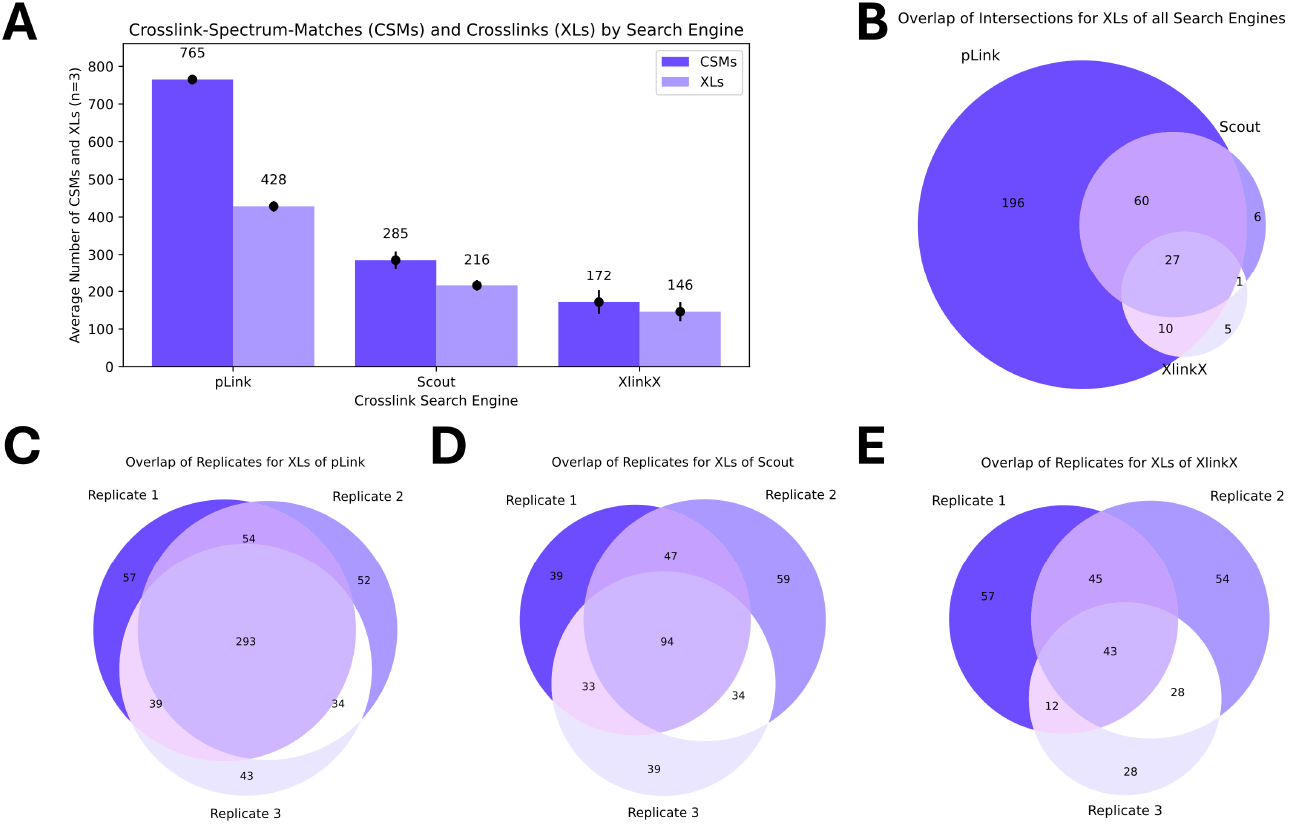
Overview of the results from pLink [39], Scout [14], and XlinkX [42] on the Cas9 crosslinking data [11] created with pyXLMS. **(A)** On average pLink identifies 765 CSMs and 428 XLs per replicate, while Scout identifies 285 CSMs and 216 XLs on average per replicate. XlinkX reports an average of 172 identified CSMs and 146 identified XLs per replicate. All results are validated for 1% estimated FDR. Error bars represent the standard deviation of three replicates (n = 3). **(B)** Overlap of identified XLs for pLink, Scout, and XlinkX when only taking XLs that are found across all three replicates by each search engine. Essentially this is a Venn diagram of the intersections of C, D, and E. **(C)** Overlap of the XLs identified by pLink across all three replicates. 293 XLs are identified in all three replicates. **(D)** Overlap of the XLs identified by Scout across all three replicates with 94 XLs which are identified in all three replicates. **(E)** Overlap of the XLs identified by XlinkX across all three replicates where 43 XLs are identified in all replicates.

**Fig. 3.**
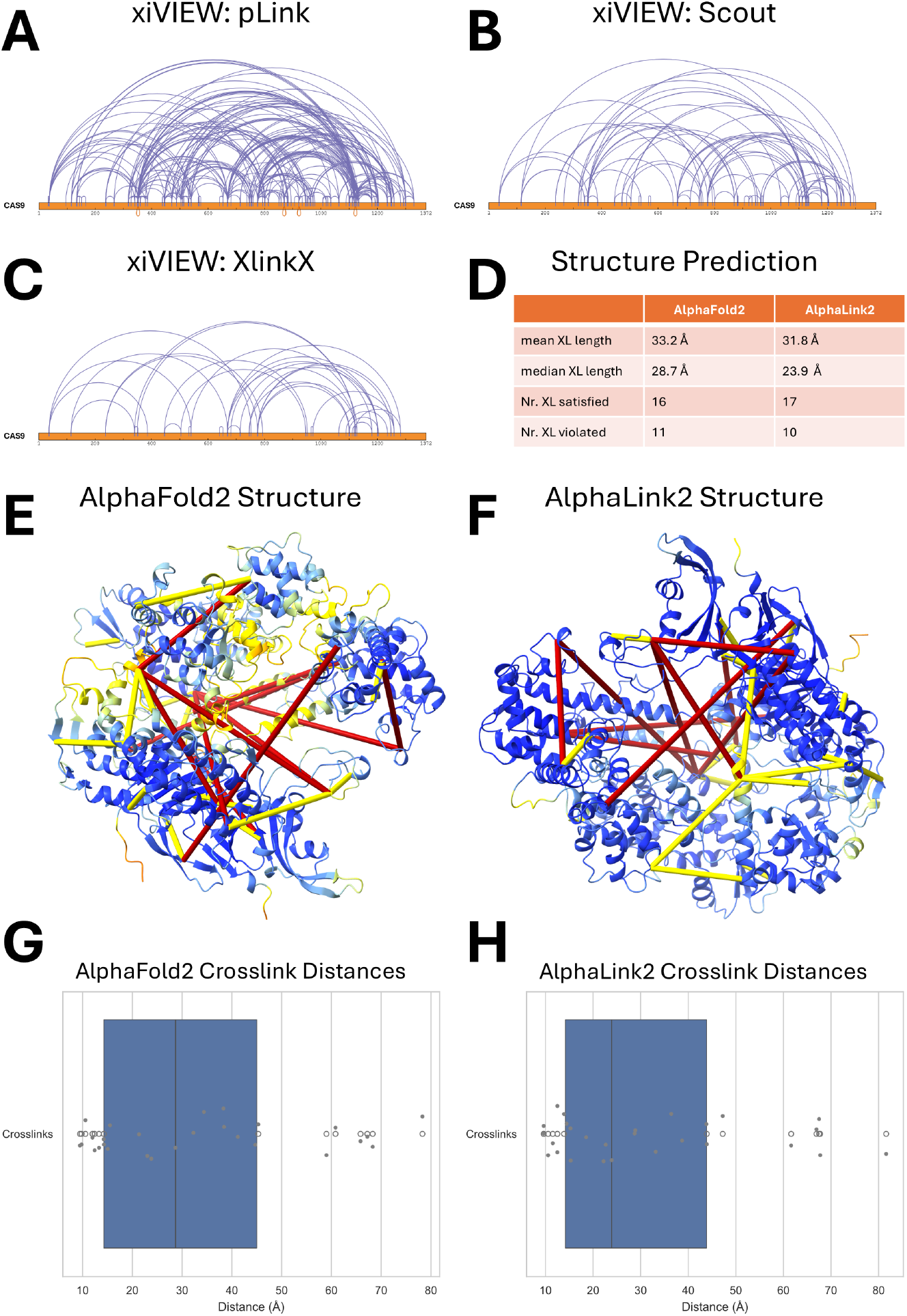
Overview of the down-stream analysis of the crosslink results. Panels A-C show all unambiguous Cas9 intra-crosslinks mapped to the sequence of Cas9 with xiVIEW [26] for crosslinks that were validated at 1% estimated FDR and identified in all three replicates by the search engine **(A)** pLink [39], **(B)** Scout [14], and **(C)** XlinkX [42]. We then took the intersection of A-C and mapped it to the AlphaFold2 [55] predicted structure of Cas9 and the structure predicted by AlphaLink2 [23] which additionally uses the crosslink constraints. Panel **(D)** shows a comparsion of the two predicted structures in terms of agreement with the crosslink results with the AlphaLink2 structure generally showing small improvements over the AlphaFold2 predicted structure. Panel **(E)** shows the AlphaFold2 predicted structure and panel **(F)** the AlphaLink2 predicted structure visualized by XMAS [50] with crosslinks that satisfy the distance threshold in yellow (16 for AlphaFold2, 17 for AlphaLink2) and crosslinks that violate the distance threshold in red (11 for AlphaFold2, 10 for AlphaLink2). The structures are colored by pLDDT using the standard AlphaFold coloring [55]. Panels G and H show the crosslink distance distributions for the **(G)** AlphaFold2 predicted structure and **(H)** the AlphaLink2 predicted structure. The AlphaLink2 predicted structure has slightly lower mean and median crosslink distances.

For this dataset pLink outperformed the other search engines not only in terms of number of identified CSMs and XLs at 1% estimated FDR with an average of 765 CSMs and 428 XLs per replicate, but also in terms of overlap between replicates with 293 XLs that are identified across all three replicates. Scout reported on average 285 CSMs and 216 XLs at an estimated FDR of 1% with an overlap of 94 XLs that were detected in all three replicates. The crosslink search engine XlinkX identified on average 172 CSMs and 146 XLs per replicate at 1% estimated FDR with 43 XLs that were detected in all three replicates. Generally the agreement between replicates is best for pLink and gradually gets more disjoint for both Scout and XlinkX. We investigated the intersections for all crosslink search engines with xiVIEW and ultimately took an intersection of all XLs that were found in all replicates by all search engines which resulted in a total of 27 XLs (see Figure 2B) that we used for further structural investigations with XMAS and AlphaLink2. Taking the intersection for down-stream analysis reduces the chance of accidentally including a false-positive crosslink identification [51].

Figure 3 shows the results of the down-stream analysis. We visualized unambiguous intra-crosslinks of Cas9 that were identified in all three replicates at 1% estimated FDR via xiVIEW [26] for the crosslink search engines pLink [39] (Figure 3A), Scout [14] (Figure 3B), and XlinkX [42] (Figure 3C), again also confirming that pLink identifies the most Cas9 intra-crosslinks. We then took the intersection of all these crosslinks with pyXLMS (27 crosslinks in total after intersection) and used them to investigate the structure of the crosslinked Cas9. In order to facilitate that, we predicted the structure of Cas9 once with AlphaFold2 [55], and once with AlphaLink2 [23] directly using the files generated with pyXLMS. In comparison to AlphaFold2, AlphaLink2 additionally uses the crosslink constraints to predict the protein structure. Subsequently, we visualized the predicted structures and crosslinks with XMAS [50] in ChimeraX [28– 30] as shown in Figure 3E and F. We examined how many of the crosslinks satisfied or violated the crosslinker specific distance constraint of DSBSO [54] which we chose as 35Å as suggested by Jiao *et al*. [56]. In total 16 crosslinks satisfied the distance constraint in the AlphaFold2 predicted structure and 17 crosslinks in the AlphaLink2 predicted structure, showing a slight improvement when using AlphaLink2. In turn, there were 11 crosslinks that violated the distance constraint in the AlphaFold2 predicted structure and 10 crosslinks in the AlphaLink2 predicted structure. Overall the mean and median crosslink distance decreased from 33.2Å to 31.8Å and from 28.7Å to 23.9Å respectively, when using AlphaLink2 compared to AlphaFold2 as shown in the crosslink distance distribution plots in Figure 3G and H. The model quality is good for both structures with an average pLDDT of 86.8 and 93.7 for AlphaFold2 and AlphaLink2 correspondingly, and the many observed over-length crosslinks are explained by the flexibility of Cas9 that inevitably cannot be easily captured in a static snapshot.

### 2.3 The pyXLMS web application provides an easy to use interface for non-programmers

We have built a web application on top of the pyXLMS python package for researchers without a bioinformatics background that offers an intuitive graphical user interface and almost all the functionality of the underlying python package without the need for any programming experience. The web application is divided into four different tabs that inherently guide researchers through the app. First, researchers can upload their results, choose a crosslink search engine or format, select a crosslinker and are then able to import their data. Secondly, users may filter their data for their desired CSMs and XLs. Thirdly, the visualization tab as depicted in Figure 4 -shows various different descriptive plots of the filtered data and users can confirm that they selected CSMs and XLs that they are interested in. Last but not least, researchers can export the data to one or more of the supported down-stream analysis tools to further investigate their data for biological significance.

**Fig. 4.**
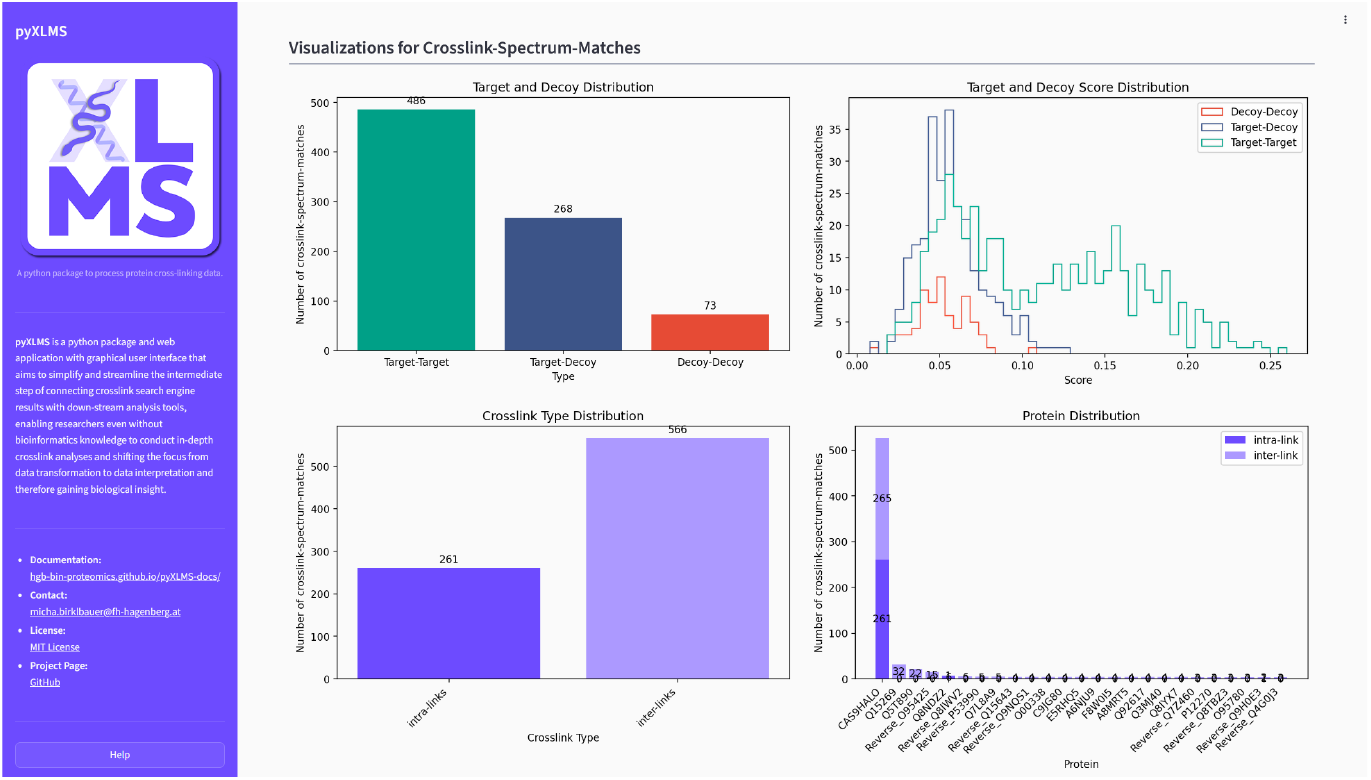
The pyXLMS web application available via https://hgb-bin-proteomics.github.io/pyXLMS-app offers a programming-free experience with an intuitive graphical user interface to transform and export crosslinking data. The here displayed visualization tab shows different plots describing the uploaded crosslinking data. In the top left an overview of the identified target-target, target-decoy, and decoy-decoy matches is shown. The top right displays a score distribution for all CSMs. The bottom left shows the distribution of identified intra- and inter-crosslinks. The bottom right shows the most frequently crosslinked proteins by both intra- and inter-crosslinks. The figure shows data of CSMs identified with Scout [14] in replicate one of the previously used dataset [11], see section 2.2.

We are hosting a free and publicly available instance of the pyXLMS web application at https://hgb-bin-proteomics.github.io/pyXLMS-app, however we also support self-hosting or running the web application locally it is cross-platform and runs on any computer that supports python and a web browser. Instructions for self-hosting, running it locally, and a user guide for navigating trough the app are available via https://hgb-bin-proteomics.github.io/pyXLMS-docs or via the Supplementary Information.

### 2.4 pyXLMS is well documented with step-by-step instructions for both programmers and non-programmers

One core focus of pyXLMS is usability and to empower all researchers in crosslinking, whether they are in the wet or dry lab, newcomers or experts in the field, or even developers of their own tools. We have compiled extensive resources that document all functionality in pyXLMS which includes not only a package reference that documents all of the python module but also an extensive user guide featuring many examples of common usage patterns - for both the python package and the web application. The user guide is available via https://hgb-bin-proteomics.github.io/pyXLMS-docs and consists of an in-depth explanation of all functionality, important background information about the crosslink search engines and down-stream analysis tools, and many examples that include step-by-step instructions. We have implemented an assortment of Jupyter notebooks to follow along, with different example data to get a hands-on experience with both pyXLMS and the supported down-stream analysis tools. The user guide is written for researchers with and without programming experience and includes instructions well beyond pyXLMS including the installation of all required tools. An exemplary excerpt of the user guide is shown in Figure 5.

**Fig. 5.**
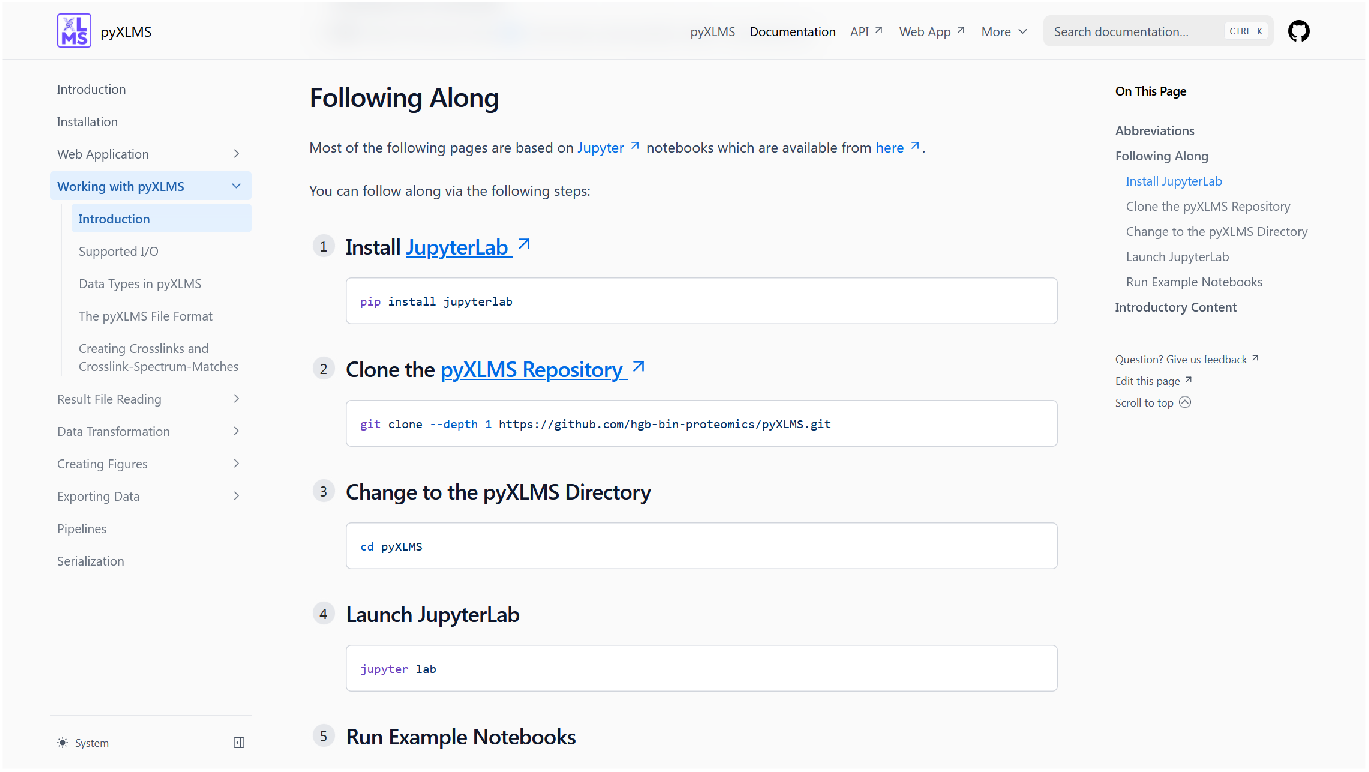
An excerpt of the pyXLMS user guide available via https://hgb-bin-proteomics.github.io/pyXLMS-docs. The pyXLMS user guide features step-by-step instructions for using the pyXLMS python package as well as the pyXLMS web application, including many common usage examples and exemplary data to try all the functionality first hand.

For developers we have created a complete python package reference at https://hgb-bin-proteomics.github.io/pyXLMS/ and have setup contribution guidelines in the pyXLMS repository.

## 3 Discussion

We here present pyXLMS, a python package and web application with graphical user interface that aims to simplify and streamline the intermediate step of connecting crosslink search engine results with down-stream analysis tools which has remained a manual, cumbersome, and error prone step in the data analysis that often requires extensive data re-shaping or transformation. This step is only possible for scientists with a strong bioinformatics background and yet is crucial for gaining biological insight from the identified crosslinks. pyXLMS is enabling scientists even without bioinformatics knowledge to conduct in-depth crosslink analyses and shifting the focus from data transformation to data interpretation. pyXLMS currently already supports input from seven different crosslink search engines, namely MaxLynx [15], MeroX [35, 36], MS Annika [13, 37, 38], pLink 2 and pLink 3 [39], Scout [14], xiSearch [40] and the validation tool xiFDR [41], XlinkX [42], as well as the mzIdentML format [31–33] of the HUPO Proteomics Standards Initiative, and a well-documented and human-readable custom tabular format. Down-stream analysis is facilitated by functionality that is directly available within pyXLMS such as aggregation, validation, annotation, filtering, and visualization of CSMs and crosslinks. Most importantly, pyXLMS enables easy export to the required data format of more than ten different down-stream analysis tools such as AlphaLink2 [23], ProXL [24, 25], xiNET [43], xiVIEW [26], xiFDR [41], XlinkDB [44–47], xlms-tools [48], PyMOL [27](via pyXlinkViewer [49]), ChimeraX [28–30](via XMAS [50]), or IMP-X-FDR [51].

We demonstrated the applicability and benefits of pyXLMS by re-analyzing a publicly available crosslink dataset [11] of the protein Cas9 crosslinked with DSBSO [54]. We show how the same data analysis workflow including down-stream analysis tools xiVIEW [26], XMAS [50], and AlphaLink2 [23] can be applied across the different crosslink search engines pLink [39], Scout [14], and XlinkX [42] using pyXLMS. The integration of pyXLMS allows us to switch between search engines and analysis tool at will, eliminating the need for manual adoption to the various data formats.

Despite our best efforts pyXLMS still comes with some limitations that are a direct result of the differences in the output formats of crosslink search engines. Many crosslink search engines do not report any kind of decoy matches which makes validation in pyXLMS or export to xiFDR [41] impossible which is why we recommend using validated results for pyXLMS. Validation within pyXLMS is currently only supported for MaxLynx [15], MS Annika [13], and xiSearch [40]. Furthermore, the different down-stream analysis tools require varying input information which might not be consistently available from all crosslink search engines. Some of this can be mitigated by functionality in pyXLMS such as annotation or by additional information that needs to be passed to pyXLMS for a successful export. Generally, the export to all downstream analysis tools should work for all crosslink search engines and input formats, with the exception of the export to xiFDR which is limited to MaxLynx, MS Annika, and xiSearch for above reasons. For safety pyXLMS makes sure before the export that all the required information is available and will otherwise throw an error.

pyXLMS is publicly available under a permissive open-source license via https://github.com/hgb-bin-proteomics/pyXLMS and we are not only welcoming but hoping for contributions by the community with the aim of supporting as many crosslink search engines and down-stream analysis tools as possible. Moreover, as long time contributors to the field of XLMS we are committed to maintain the project and drive development forward. We think that pyXLMS is an important step in making XLMS data analysis more accessible to everyone and believe that it has the potential to become a foundational building block in future crosslinking tools.

## 4 Methods

### 4.1 Implementation of pyXLMS

pyXLMS is implemented in python using a functional programming approach and divided into currently four submodules: (1) *parser* handles reading of all input files from the different crosslink search engines and data formats. For reading of tabular data the python package *pandas* [57, 58] is used, for reading of mzIdentML files we make use of *pyteomics* [59, 60]. All data is parsed into python native data structures for performance and serializability, this is further described in the pyXLMS documentation. (2) The submodule *transform* handles all data transformation steps including filtering, annotation, and validation. For reading FASTA files we use *biopython* [61], all other functionality is natively implemented. For validation we implemented the FDR estimation algorithm as used by MS Annika [13] as well as the formula suggested by Fischer and Rappsilber for directional crosslinks [41]. (3) The *plotting* submodule implements all visualization functionality and mainly uses the python package *matplotlib* [62]. (4) The *exporter* submodule handles exporting CSMs and XLs to the formats required by the supported down-stream analysis tools. The python package *biopandas* [63] is used for reading PDB files and *biopython* [61] is used for sequence alignment. We use *lxml* for XML schema validation of ProXL [24, 25] exports.

The web application is implemented in *streamlit*. For documentation we use *sphinx* and *nextra*. All python code is linted with *ruff*, type checked with *pyright*, and tested with *pytest*. We currently have 584 tests covering the codebase of pyXLMS, that includes tests for parsing of the result files of all supported crosslink search engines. We tested result files from MaxLynx as part of MaxQuant version 2.6.2.0 [15, 34], MeroX version 2.0.1.4 [35, 36], MS Annika version 3.0.7 [13], pLink version 2.3.11 and version 3.0.17 [39], Scout version 1.6.3 (released by Diego Borges, https://github.com/diogobor/Scout/) and version 2.0.0 beta (released by The Liu Lab, https://github.com/theliulab/Scout/) [14], xiSearch version 1.7.6.7 [40], xiFDR version 2.2.1 [41], XlinkX as part of Thermo Fisher Scientific’s Proteome Discoverer versions 3.1.0.638 and 3.2.0.450 [42], as well as the mzIdentML format version 1.2 [32].

### 4.2 Data acquisition

All used mass spectrometry proteomics data was retrieved from the ProteomeXchange consortium (http://proteomecentral.proteomexchange.org) [64] via the PRIDE partner repository [53] with the dataset identifier PXD061173 [11]. In short, the authors prepared purified recombinant Cas9 from *S. pyogenes* fused with a Halo-tag and crosslinked it with DSBSO. The crosslinked Cas9 samples were spiked in a 1:80 (w/w, 20 µg Cas9 + 1.6 mg HeLa) ratio into a HeLa protein digest, followed by DBCO bead-based enrichment. Samples were then analyzed using a Vanquish Neo UHPLC system and an Orbitrap Eclipse Tribrid mass spectrometer (both Thermo Fisher Scientific). A detailed description of all sample preparation, enrichment, and data acquisition steps is given in the authors’ original publication [11].

### 4.3 Data analysis

RAW files were directly searched with the crosslink search engines pLink [39] (version 3.0.17), Scout [14] (version 2.0.0 beta, retrieved from The Liu Lab https://github.com/theliulab/Scout/), and XlinkX [42] (version as part of Thermo Fisher Scientific’s Proteome Discoverer version 3.2.0.450) using the full human SwissProt reference proteome (n = 20 328, UniProt Proteome ID UP000005640, retrieved 03. November 2025) and the sequence of Cas9 as the protein database. Search settings were set analogous for all search engines using digestion enzyme trypsin with a maximum of three missed cleavages allowed, a minimum peptide length of six amino acids and a maximum peptide length of 30 amino acids. Carbamidomethylation of cysteine was set as a fixed modification and oxidation of methionine was set as a variable modification for all peptides. For non-crosslinked peptides we also considered the amidated, hydrolyzed and tris forms of DSBSO [54] as variable modifications for all lysine residues. The crosslinker parameter was set to DSBSO with reactions to lysine and the protein n-terminus allowed. The precursor mass tolerance was set to 5 ppm and the fragment mass tolerance to 10 ppm. For XlinkX the “Cleavable MS2” workflow was used. Results were validated for 1% estimated FDR using the in-built FDR estimation algorithms of all crosslink search engines and setting the target FDR to 1% at all possible levels. All other crosslink search engine parameters were left to their defaults.

The validated crosslink search engine results were then directly loaded with pyXLMS. Subsequently, re-annotation of associated proteins and protein crosslink positions using the *transform*.*reannotate positions()* function of pyXLMS was performed to have consistent annotation accross all search engine results. Any non-unique CSMs and crosslinks were filtered out using the pyXLMS function *transform*.*unique()* via grouping by protein crosslink position. Crosslink intersections were created using the pyXLMS intersection and Venn diagram functionalities, and were finally exported to xiVIEW [26], XMAS [50], and AlphaLink2 [23] format using the pyXLMS exporter submodule.

Results were then visualized with xiVIEW, modelled with AlphaLink2 (version git@d54bcbf), and analyzed with XMAS (version 1.1.3) in ChimeraX [28–30] (version 1.10.1) - all directly with the outputs created by pyXLMS. For the comparison to AlphaFold2 we predicted the structure of Cas9 with ColabFold [65] using the AlphaFold2 [55] notebook with MMseqs2 support [66] (notebook version 1.5.5). The sequence of Cas9 was used as input and all other ColabFold parameters were left at their default values. The rank 1 structure was used for comparison with the AlphaLink2 predicted structure.

## 5 Data availability

All used mass spectrometry proteomics data is available from the ProteomeXchange consortium (http://proteomecentral.proteomexchange.org) [64] via the PRIDE partner repository [53] with the dataset identifier PXD061173 [11]. Result files of the re-analysis, structure prediction, scripts and figure code used for this manuscript are available via the GitHub repository https://github.com/hgb-bin-proteomics/pyXLMS-manuscript.

## 6 Code availability

The python package is available from the Python Package Index (PyPI) via https://pypi.org/project/pyXLMS/ and can easily installed via “pip install pyxlms”. The source code for pyXLMS is available via the GitHub repository https://github.com/hgb-bin-proteomics/pyXLMS. The web application is hosted by the University of Applied Sciences Upper Austria and accessible free of charge via https://hgb-bin-proteomics.github.io/pyXLMS-app. Source code for the web application is part of the pyXLMS git repository. Documentation for the python package is available at https://hgb-bin-proteomics.github.io/pyXLMS/. A comprehensive user guide and documentation for both the python package and the web application is available at https://hgb-bin-proteomics.github.io/pyXLMS-docs and https://pyxlms.dev. An overview of all published pyXLMS resources is also given in the Supplementary Information.

## 7 Author contributions

M.J.B. conceptualized, implemented, tested and documented pyXLMS, performed data analysis and visualisation, and wrote the manuscript. L.M.B. and S.K. implemented, documented and tested pyXLMS. F.M. and M.M. conceptualized and tested pyXLMS, provided data, and helped with data analysis. K.M., S. W., and V.D. supervised the project. All authors revised the manuscript and have given approval to the final version of the manuscript.

## 9 Ethics approval and consent to participate

Not applicable.

## 10 Consent for publication

Not applicable.

## 11 Materials availability

Not applicable.

**Supplementary information**

The following supplementary information is provided:

- Supplementary Information.pdf:

1. Supplementary Section 1: pyXLMS Quick Links.
2. Supplementary Section 2: pyXLMS Web App User Manual.

## Acknowledgements

This work was supported by the F&E Infrastruk- turförderung 4. Ausschreibung 2022/01 (AT-SCP, https://projekte.ffg.at/projekt/4795911, accessed on 24 November 2025) of the Austrian Research Promotion Agency (FFG). This work was further funded by the project LS20-079 of the Vienna Science and Technology Fund and the project P35045-B of the Austrian Science Fund (FWF, Grant DOI 10.55776/P35045) and the the ESPRIT program project number ESP 566 (Grant-DOI 10.55776/ESP566). Additionally, this work was supported by the MSCA-ITN-2020 PROTrEIN project that has received funding from the European Union’s Horizon 2020 programme under the Marie Sklodowska-Curie grant agreement No. 956148. Furthermore, we thank Martina A. Höllwarth, Eva Rauch, and Dennis Dannecker who gave feedback and tested pyXLMS. We also thank Melanie E. Birklbauer who designed the logo for pyXLMS.

This research was funded in whole, or in part, by the Austrian Science Fund (FWF). For the purpose of open access, the author has applied a CC BY public copyright licence to any Author Accepted Manuscript version arising from this submission.

