## Supplementary Information for "Unified down-stream analysis of crosslinking mass spectrometry results with pyXLMS"

Micha J. Birklbauer[1,2]*, Louise M. Buur[1,2], Sabrina Kaser[1], Fränze Müller[3], Manuel Matzinger[3], Karl Mechtler[3,4,5], Stephan Winkler[1,2], Viktoria Dorfer[1]*

[1] Bioinformatics Research Group, University of Applied Sciences Upper Austria, Softwarepark 11, Hagenberg, 4232, Upper Austria, Austria.
[2] Institute for Symbolic Artificial Intelligence, Johannes Kepler University Linz, Altenberger Straße 69, Linz, 4040, Upper Austria, Austria.
[3] Institute of Molecular Pathology (IMP), Vienna BioCenter (VBC), Campus-Vienna-Biocenter 1, Vienna, 1030, Vienna, Austria.
[4] Institute of Molecular Biotechnology (IMBA), Austrian Academy of Sciences, Vienna BioCenter (VBC), Dr. Bohr-Gasse 3, Vienna, 1030, Vienna, Austria.
[5] Gregor Mendel Institute (GMI), Austrian Academy of Sciences, Vienna BioCenter (VBC), Dr. Bohr-Gasse 3, Vienna, 1030, Vienna, Austria.

### Table of Contents

**Table of Contents** 2
**Supplementary Section 1 - pyXLMS Quick Links** 3
**Supplementary Section 2 - pyXLMS Web App User Manual** 4
Loading Data 4
File Selection 4
Selecting Crosslink Search Engine or File Format 5
Selecting Crosslink Reagent 5
Reading the Results 5
[Optional] Pre-Processing Results 6
[Optionally] Apply One or More Pre-Processing Steps 6
Reading the Results 7
Inspecting Your Results 8
Filtering Results 9
Filter Options 9
Filter by Protein Accession 9
Filter by Crosslink Type 9
Filter by Target-Decoy Type 10
Inspect Your Filtered Results 10
Visualizing Results 11
Exporting Results 13
Exporting Your Results 13
Select the Down-Stream Analysis Tool/Export Format 13
Verify Your Pre-Processing and Filtering 14
Supply Any Additionally Required Data 14
Create the Export 15
Download the Exported File(s) 15
Use Your Exported File(s) with the Selected Down-Stream Analysis Tool 15
Getting Help 16

### Supplementary Section 1 - pyXLMS Quick Links

Here is a quick overview of the most important pyXLMS resources:

- Web Application: https://hgb-bin-proteomics.github.io/pyXLMS-app
- User Guide: https://hgb-bin-proteomics.github.io/pyXLMS-docs
- GitHub Repository: https://github.com/hgb-bin-proteomics/pyXLMS

Here is a complete overview of all available pyXLMS resources:

- https://github.com/hgb-bin-proteomics/pyXLMS: pyXLMS git repository, source code, and main page
- https://hgb-bin-proteomics.github.io/pyXLMS-app: publicly hosted instance of the pyXLMS web application that is free to use for everyone
- https://hgb-bin-proteomics.github.io/pyXLMS-docs: User guide with in-depth step-by-step instructions for using both the python package and the web application
- https://hgb-bin-proteomics.github.io/pyXLMS/: pyXLMS python package documentation
- https://github.com/hgb-bin-proteomics/pyXLMS/tree/master/gui: source code of the pyXLMS web application
- https://github.com/hgb-bin-proteomics/pyXLMS-app: information about our publicly hosted instance of the pyXLMS web application
- https://github.com/hgb-bin-proteomics/pyXLMS-docs: source code of the user guide website
- https://github.com/hgb-bin-proteomics/pyXLMS-manuscript: source code for all analyses of the manuscript

### Supplementary Section 2 - pyXLMS Web App User Manual

This guide can also be found at https://pyxlms.dev/docs/webapp.

#### Loading Data

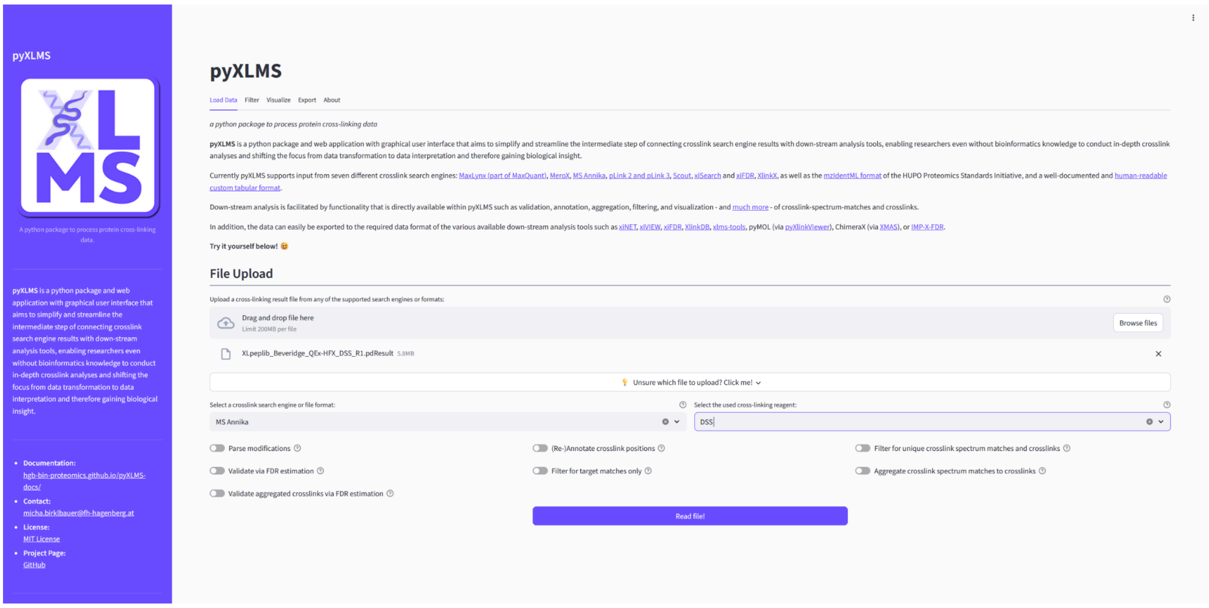


##### File Selection

The first step when using the pyXLMS web application is that you need to upload one or more result files containing crosslink-spectrum-matches (CSMs) or crosslinks/residue pairs (XLs) in the Load Data tab. You can find a list of supported crosslink search engines and input formats here.

We also provide some example files via the pyXLMS GitHub repository that you can try!

**TIP**

*Starting from pyXLMS v1.6.1 the web application also supports input of multiple result files, e.g. if you have CSMs and XLs from the same run, result files of different fractions of your sample, or replicates.*

#### Selecting Crosslink Search Engine or File Format

Next you need to select the correct crosslink search engine or file format from the dropdown menu. This should simply be the crosslink search engine that you used for crosslink identification with your mass spectrometry data, or a generic format like mzIdentML.

#### Selecting Crosslink Reagent

Then you need to select the crosslinking reagent that was used in your experiment from the next dropdown menu. If your reagent is not in the list you can select Custom and two additional input fields will pop up where you can specify a crosslinker name and the monoisotopic delta mass of the crosslink modification.

#### Reading the Results

If you don't want to employ any of the additional and optional pre-processing steps (see next section) you can read your results now by clicking on the Read file(s)! button!

### [Optional] Pre-Processing Results

pyXLMS offers several different pre-processing steps as outlined below.

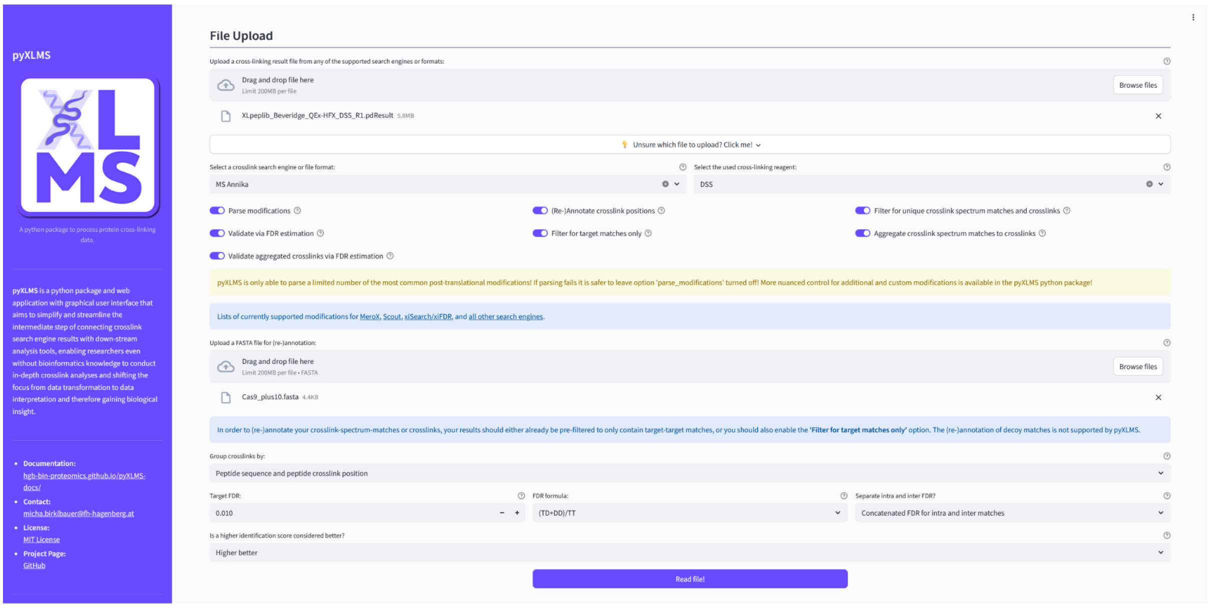


#### [Optionally] Apply One or More Pre-Processing Steps

- **Parse Modifications**

  Enabling this option will try to parse post-translation-modifications (PTMs) of your CSMs from your result files.

  **NOTE**

  *Please note that the pyXLMS web app only supports a limited set of the most common PTMs! If parsing fails it is better to leave this option turned off and use the pyXLMS python package directly which gives a more nuanced control over modifications!*

- **Re-Annotate Crosslink Positions**

  This option will allow you to upload a FASTA file to re-annotate the protein crosslink positions of your CSMs and XLs based on the given FASTA file. This might be useful if the annotation is missing because it was not given by the crosslink search engine, or if the crosslink search engine annotated something wrongly.

- **Filtering for Unique CSMs and XLs**
  If turned on this option will filter out any non-unique CSMs and XLs from your results. A crosslink is considered unique if its sequence and peptide crosslink positions are unique (option 1) or if its protein crosslink positions are unique (option 2) - these options are controlled via the Group crosslinks by selector.
- **Validation via FDR Estimation**
  Enabling this option will validate your results for a given target false-discover-rate (FDR) via FDR estimation and will only keep CSMs and XLs that fall within the target FDR.

  **IMPORTANT**
  *We recommend uploading results that are already validated in the crosslink search engine of your choice. Validation in pyXLMS uses a generic target-decoy approach with a single score that might not yield the same results as the search engine specific validation!*
- **Filtering for Target Matches**
  If applied this option will filter out any CSMs and XLs that are not target-target matches.
- **Aggregation of CSMs to XLs**
  Whether or not your CSMs (if there are any) should be aggregated to XLs.
- **Validation of Aggregated XLs**
  Controls if your aggregated XLs from the previous pre-processing step should also be validated via FDR estimation. Shares the same settings as Validation via FDR Estimation.

#### Reading the Results

You can read your results now by clicking on the Read file(s)! button!

### Inspecting Your Results

Depending on the result file(s) you uploaded and the selected pre-processing steps you will see the read CSMs and/or XLs and/or aggregated XLs. The web app will both display the read data itself as well as a short summary statistic. You will also be able to directly download the data in different formats.

##### Read Crosslink-Spectrum-Matches

**Read 699 crosslink-spectrum-matches:**

| | Completeness | Alpha Peptide | Alpha Peptide Modifications | Alpha Peptide Crosslink Position | Alpha Proteins | Alpha Proteins Crosslink Positions | Alpha Proteins Peptide Positions | Alpha Score | Alpha Decoy | Beta Peptide | Beta Peptide Modifications | Beta Peptide Crosslink Position | Beta Proteins | Beta Proteins Cr |
|---|---|---|---|---|---|---|---|---|---|---|---|---|---|---|
| 0 | full | GQKNSR | (3:[DSS\|138.06808]) | 3 | Cas9 | 779 | 777 | 119.8255 | ☐ | GQKNSR | (3:[DSS\|138.06808]) | 3 | Cas9 | 779 |
| 1 | full | SDKNR | (3:[DSS\|138.06808]) | 3 | Cas9 | 866 | 864 | 114.4283 | ☐ | SDKNR | (3:[DSS\|138.06808]) | 3 | Cas9 | 866 |
| 2 | full | DKQSGK | (2:[DSS\|138.06808]) | 2 | Cas9 | 677 | 676 | 200.976 | ☐ | DKQSGK | (2:[DSS\|138.06808]) | 2 | Cas9 | 677 |
| 3 | full | DKQSGK | (2:[DSS\|138.06808]) | 2 | Cas9 | 677 | 676 | 94.4706 | ☐ | HSIKK | (4:[DSS\|138.06808]) | 4 | Cas9 | 48 |
| 4 | full | VPSKK | (4:[DSS\|138.06808]) | 4 | Cas9 | 34 | 31 | 110.4822 | ☐ | VPSKK | (4:[DSS\|138.06808]) | 4 | Cas9 | 34 |
| 5 | full | GQKNSR | (3:[DSS\|138.06808]) | 3 | Cas9 | 779 | 777 | 117.5004 | ☐ | GYKEVK | (3:[DSS\|138.06808]) | 3 | Cas9 | 1192 |
| 6 | full | KVTVK | (1:[DSS\|138.06808]) | 1 | Cas9 | 562 | 562 | 152.9711 | ☐ | KVTVK | (1:[DSS\|138.06808]) | 1 | Cas9 | 562 |
| 7 | full | EKIEK | (2:[DSS\|138.06808]) | 2 | Cas9 | 443 | 442 | 122.4293 | ☐ | KVTVK | (1:[DSS\|138.06808]) | 1 | Cas9 | 562 |
| 8 | full | LSKSR | (3:[DSS\|138.06808]) | 3 | Cas9 | 222 | 220 | 113.5902 | ☐ | LSKSR | (3:[DSS\|138.06808]) | 3 | Cas9 | 222 |
| 9 | full | EKIEK | (2:[DSS\|138.06808]) | 2 | Cas9 | 443 | 442 | 163.1031 | ☐ | EKIEK | (2:[DSS\|138.06808]) | 2 | Cas9 | 443 |

**Summary Statistics:**

| Number of CSMs | Number of unique CSMs | Number of intra CSMs | Number of inter CSMs | Number of target-target CSMs | Number of target-decoy CSMs | Number of decoy-decoy CSMs | Minimum CSM score | Maximum CSM score |
|---|---|---|---|---|---|---|---|---|
| 699 | 699 | 696 | 3 | 699 | 0 | 0 | 34.1885 | 452.9862 |

Download crosslink-spectrum-matches as .csv! | Download crosslink-spectrum-matches as .xlsx! | Download crosslink-spectrum-matches as .json!

➡ Exemplary depiction of read CSMs.

##### Read Crosslinks

**Read 224 crosslinks:**

| | Completeness | Alpha Peptide | Alpha Peptide Crosslink Position | Alpha Proteins | Alpha Proteins Crosslink Positions | Alpha Decoy | Beta Peptide | Beta Peptide Crosslink Position | Beta Proteins | Beta Proteins Crosslink Positions | Beta Decoy | Crosslink Type | Crosslink Score |
|---|---|---|---|---|---|---|---|---|---|---|---|---|---|
| 0 | full | GQKNSR | 3 | Cas9 | 779 | ☐ | GQKNSR | 3 | Cas9 | 779 | ☐ | intra | 119.8255 |
| 1 | full | SDKNR | 3 | Cas9 | 866 | ☐ | SDKNR | 3 | Cas9 | 866 | ☐ | intra | 114.4283 |
| 2 | full | DKQSGK | 2 | Cas9 | 677 | ☐ | DKQSGK | 2 | Cas9 | 677 | ☐ | intra | 200.976 |
| 3 | full | DKQSGK | 2 | Cas9 | 677 | ☐ | HSIKK | 4 | Cas9 | 48 | ☐ | intra | 94.4706 |
| 4 | full | VPSKK | 4 | Cas9 | 34 | ☐ | VPSKK | 4 | Cas9 | 34 | ☐ | intra | 110.4822 |
| 5 | full | GQKNSR | 3 | Cas9 | 779 | ☐ | GYKEVK | 3 | Cas9 | 1192 | ☐ | intra | 117.5004 |
| 6 | full | KVTVK | 1 | Cas9 | 562 | ☐ | KVTVK | 1 | Cas9 | 562 | ☐ | intra | 152.9711 |
| 7 | full | EKIEK | 2 | Cas9 | 443 | ☐ | KVTVK | 1 | Cas9 | 562 | ☐ | intra | 122.4293 |
| 8 | full | LSKSR | 3 | Cas9 | 222 | ☐ | LSKSR | 3 | Cas9 | 222 | ☐ | intra | 116.8895 |
| 9 | full | EKIEK | 2 | Cas9 | 443 | ☐ | EKIEK | 2 | Cas9 | 443 | ☐ | intra | 163.103 |

**Summary Statistics:**

| Number of crosslinks | Number of unique crosslinks by peptide | Number of unique crosslinks by protein | Number of intra crosslinks | Number of inter crosslinks | Number of target-target crosslinks | Number of target-decoy crosslinks | Number of decoy-decoy crosslinks | Minimum crosslink score | Maximum crosslink score |
|---|---|---|---|---|---|---|---|---|---|
| 224 | 224 | 224 | 223 | 1 | 224 | 0 | 0 | 52.9242 | 452.9862 |

Download crosslinks as .csv! | Download crosslinks as .xlsx! | Download crosslinks as .json!

➡ Exemplary depiction of read XLs.

##### Aggregated Crosslinks

**Aggregated 224 crosslinks:**

| | Completeness | Alpha Peptide | Alpha Peptide Crosslink Position | Alpha Proteins | Alpha Proteins Crosslink Positions | Alpha Decoy | Beta Peptide | Beta Peptide Crosslink Position | Beta Proteins | Beta Proteins Crosslink Positions | Beta Decoy | Crosslink Type | Crosslink Score |
|---|---|---|---|---|---|---|---|---|---|---|---|---|---|
| 0 | full | GQKNSR | 3 | Cas9 | 779 | ☐ | GQKNSR | 3 | Cas9 | 779 | ☐ | intra | 119.8255 |
| 1 | full | SDKNR | 3 | Cas9 | 866 | ☐ | SDKNR | 3 | Cas9 | 866 | ☐ | intra | 114.4283 |
| 2 | full | DKQSGK | 2 | Cas9 | 677 | ☐ | DKQSGK | 2 | Cas9 | 677 | ☐ | intra | 200.976 |
| 3 | full | DKQSGK | 2 | Cas9 | 677 | ☐ | HSIKK | 4 | Cas9 | 48 | ☐ | intra | 94.4706 |
| 4 | full | VPSKK | 4 | Cas9 | 34 | ☐ | VPSKK | 4 | Cas9 | 34 | ☐ | intra | 110.4822 |
| 5 | full | GQKNSR | 3 | Cas9 | 779 | ☐ | GYKEVK | 3 | Cas9 | 1192 | ☐ | intra | 117.5004 |
| 6 | full | KVTVK | 1 | Cas9 | 562 | ☐ | KVTVK | 1 | Cas9 | 562 | ☐ | intra | 152.9711 |
| 7 | full | EKIEK | 2 | Cas9 | 443 | ☐ | KVTVK | 1 | Cas9 | 562 | ☐ | intra | 122.4293 |
| 8 | full | LSKSR | 3 | Cas9 | 222 | ☐ | LSKSR | 3 | Cas9 | 222 | ☐ | intra | 116.8895 |
| 9 | full | EKIEK | 2 | Cas9 | 443 | ☐ | EKIEK | 2 | Cas9 | 443 | ☐ | intra | 163.103 |

**Summary Statistics:**

| Number of crosslinks | Number of unique crosslinks by peptide | Number of unique crosslinks by protein | Number of intra crosslinks | Number of inter crosslinks | Number of target-target crosslinks | Number of target-decoy crosslinks | Number of decoy-decoy crosslinks | Minimum crosslink score | Maximum crosslink score |
|---|---|---|---|---|---|---|---|---|---|
| 224 | 224 | 224 | 223 | 1 | 224 | 0 | 0 | 52.9242 | 452.9862 |

Download aggregated crosslinks as .csv! | Download aggregated crosslinks as .xlsx! | Download aggregated crosslinks as .json!

➡ Exemplary depiction of aggregated XLs created from the read CSMs.

### Filtering Results

The Filter tab offers some additional filtering options:

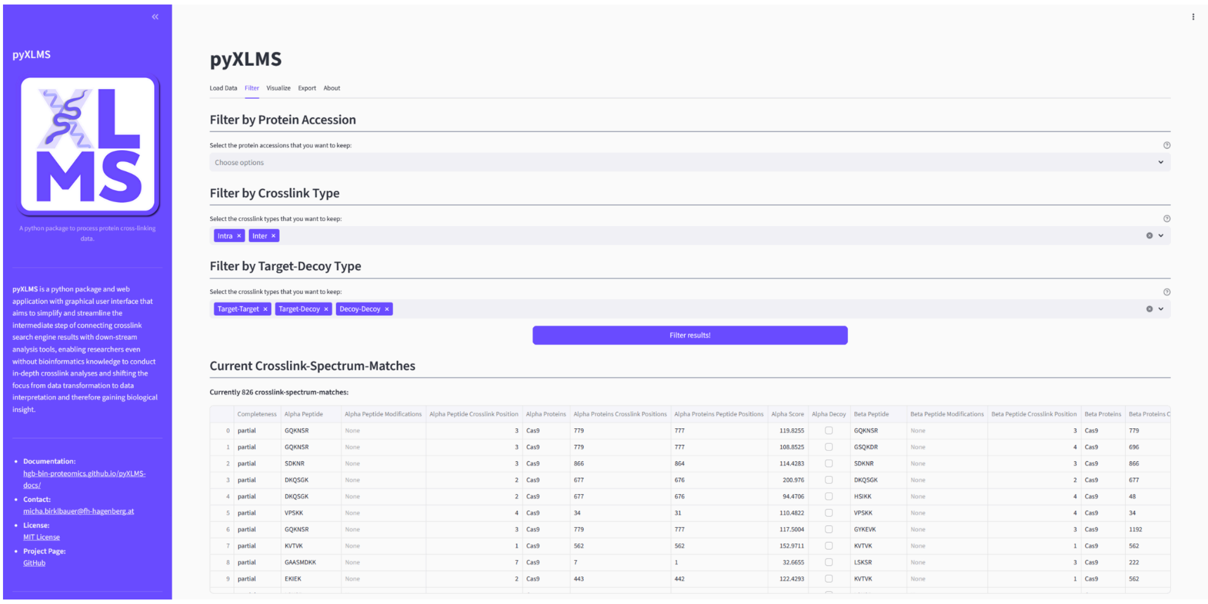


### Filter Options

#### Filter by Protein Accession

Select your proteins of interest from the dropdown menu. Any CSM or XL that does not have at least one peptide corresponding to one of the selected proteins will be filtered out. Leaving this option blank will not apply the filter.

#### Filter by Crosslink Type

Select which type of crosslinks you are interested in. Leaving this option blank will not apply the filter.

##### Filter by Target-Decoy Type

Select which database match types you are interested in. Option Target-Decoy corresponds to both "Target-Decoy" and "Decoy-Target" matches. Leaving this option blank will not apply the filter.

**IMPORTANT**

*Please note that this filter will remove any CSMs and XLs that have missing target-decoy labels!*

#### Inspect Your Filtered Results

After hitting the Filter results! button the data at the bottom of the page will update and display your current CSMs and/or XLs and/or aggregated XLs after the filtering.

### Visualizing Results

The Visualize tab will show some basic plots for your CSMs and/or XLs and/or aggregated XLs.

**NOTE**

*Please note that the visualizations are heavily influenced by your pre-processing and filtering steps! E.g. filtering for target matches will obviously mean that Target and Decoy plots will only show target-target matches!*

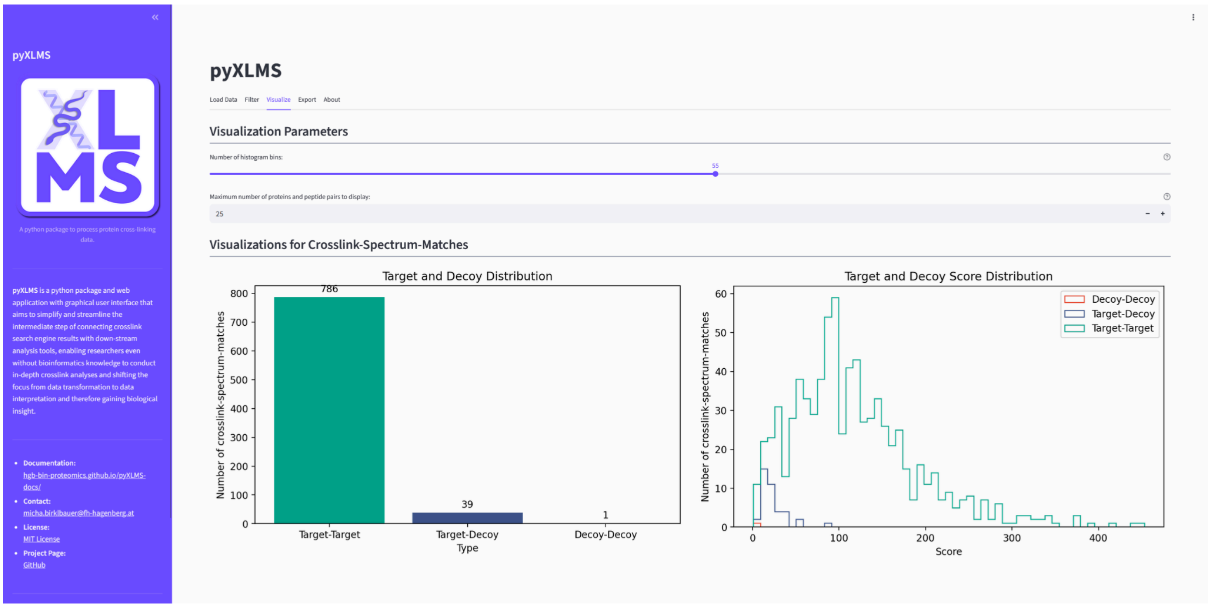


- The Target and Decoy Distribution plot will show the number of "Target-Target", "Target-Decoy" and "Decoy-Decoy" matches in your results.
- The Target and Decoy Score Distribution plot will show the score distribution of "Target-Target", "Target-Decoy" and "Decoy-Decoy" matches in your results.

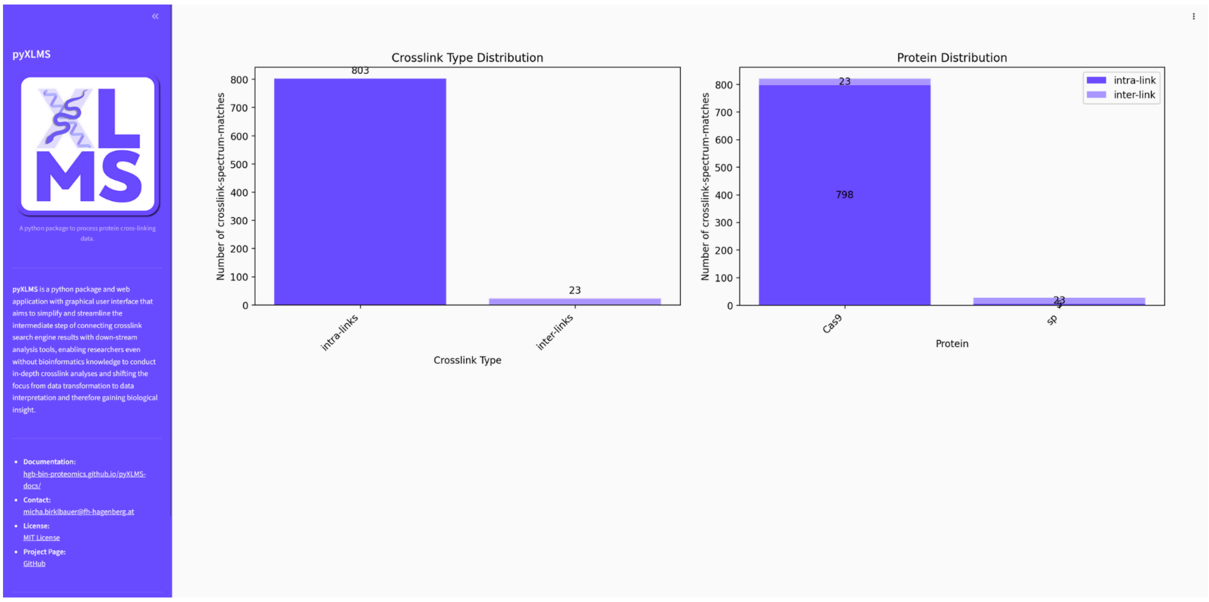


- The Crosslink Type Distribution plot will show the number of intra- and inter-links in your results.
- The Protein Distribution plot will show the top *n* proteins with the most crosslink evidence in terms of number of CSMs/XLs. It additionally shows the number of intra- and inter-links per protein. The *n* parameter is controlled via the Visualization Parameters section at the top of the page.

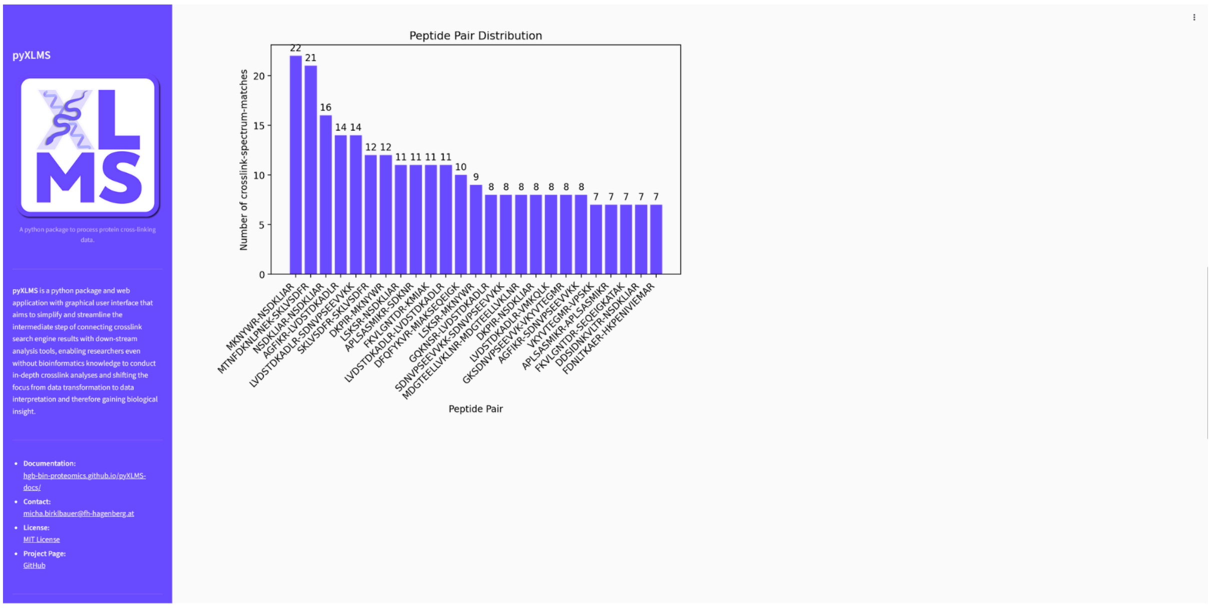


- The Peptide Pair Distribution plot is only available for CSMs and shows the most frequently observed peptide pairs in your CSMs.

### Exporting Results

The Export tab allows you to export your current selection of CSMs and/or XLs and/or aggregated XLs to one of the supported down-stream analysis tools or export formats.

**IMPORTANT**

*Please note that all exporters check if the required information to create a successful export is available at runtime! If key information is missing, an error will be raised!*

### Exporting Your Results

#### Select the Down-Stream Analysis Tool/Export Format

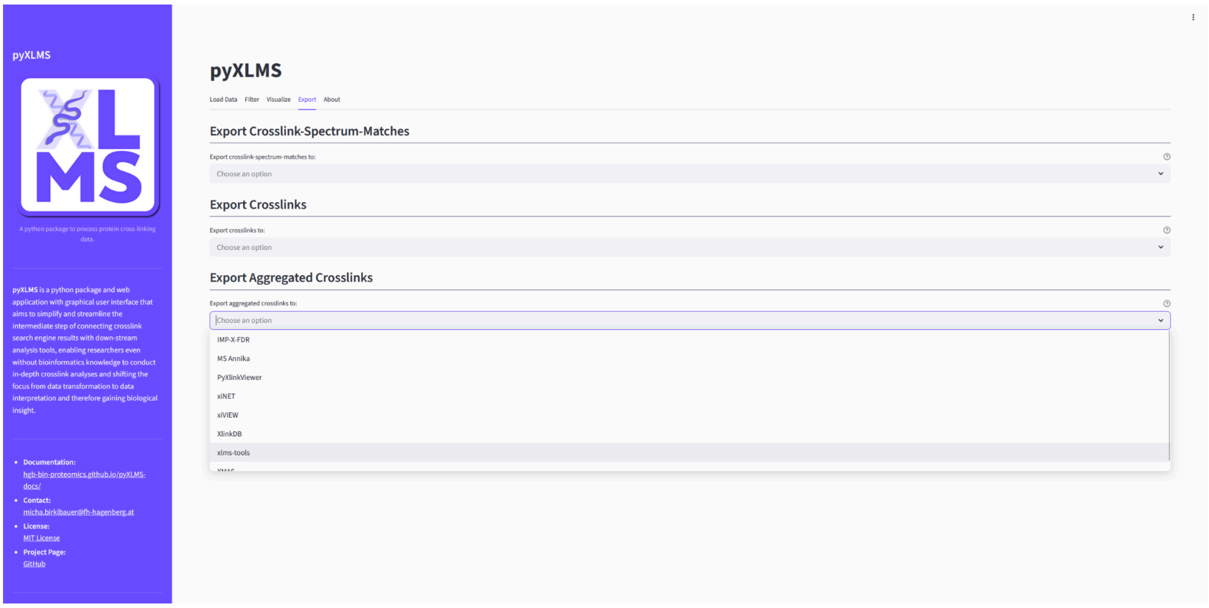


Select the down-stream analysis tool or export format that you want to get.

#### Verify Your Pre-Processing and Filtering

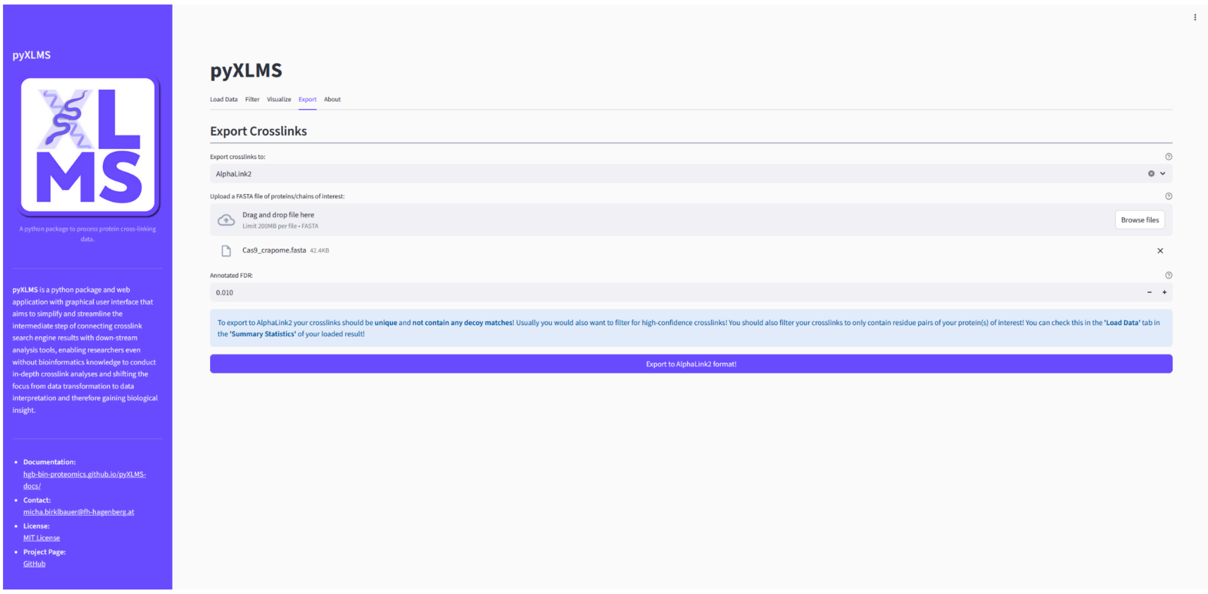


After selecting an export option a blue callout will appear with recommendations on how your results should be pre-processed and filtered for the export to work correctly. This is not only important for the export but subsequent down-stream analysis, e.g. down-stream analysis tools might not work correctly if your data is not sufficiently pre-processed and filtered, even if the export worked flawlessly.

**WARNING**

*Please note that the exporters do not check which pre-processing steps have been carried out beforehand! Some down-stream analysis tools might require validated results, some might require raw results, some might require target matches only, some might require target and decoy matches, etc. Which filtering steps are applied before export are up to you - the user - and your own responsibility. Please read the instructions of the down-stream analysis tool you want to use and get familiar with it and the required data before exporting! The exporter tutorial pages also give a short overview of recommended pre-processing steps!*

#### Supply Any Additionally Required Data

Some exporters need additional data to work, for example a FASTA file or other not yet known meta information. Please fill in this information.

#### Create the Export

Click the Export to * button to run the exporter.

#### Download the Exported File(s)

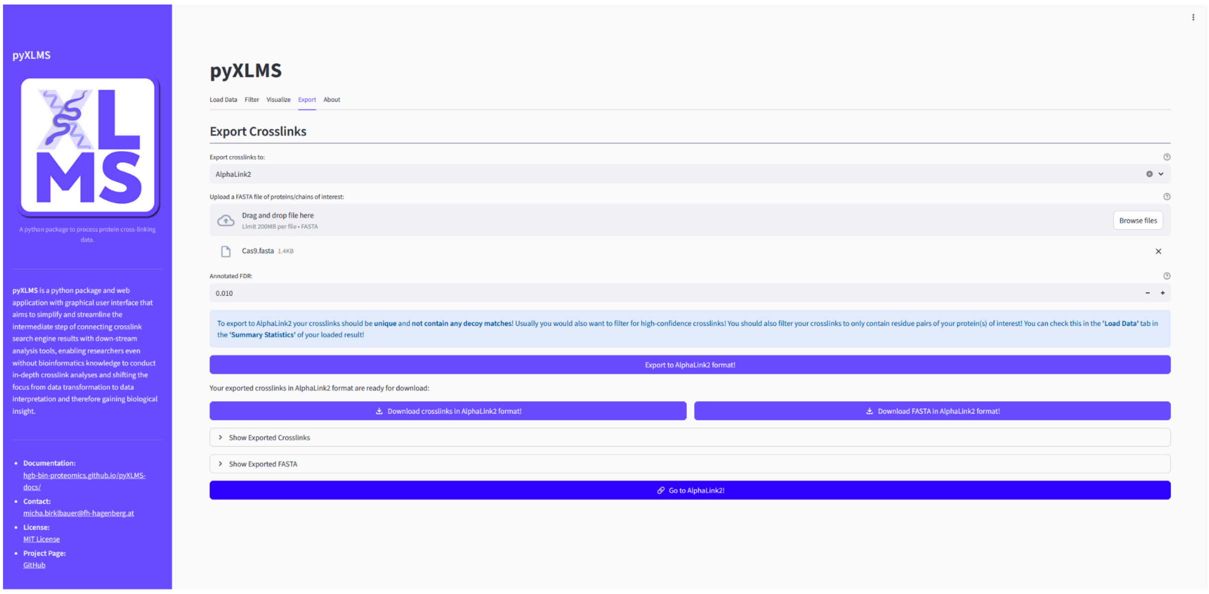


If the export finished successfully you will get the option to download the exported file(s). Sometimes also some additional information about the export is displayed.

#### Use Your Exported File(s) with the Selected Down-Stream Analysis Tool

For convenience the web app will also display a button that takes you directly to the selected down-stream analysis tool!

### Getting Help

The About tab links some useful pages on where to learn more about pyXLMS.

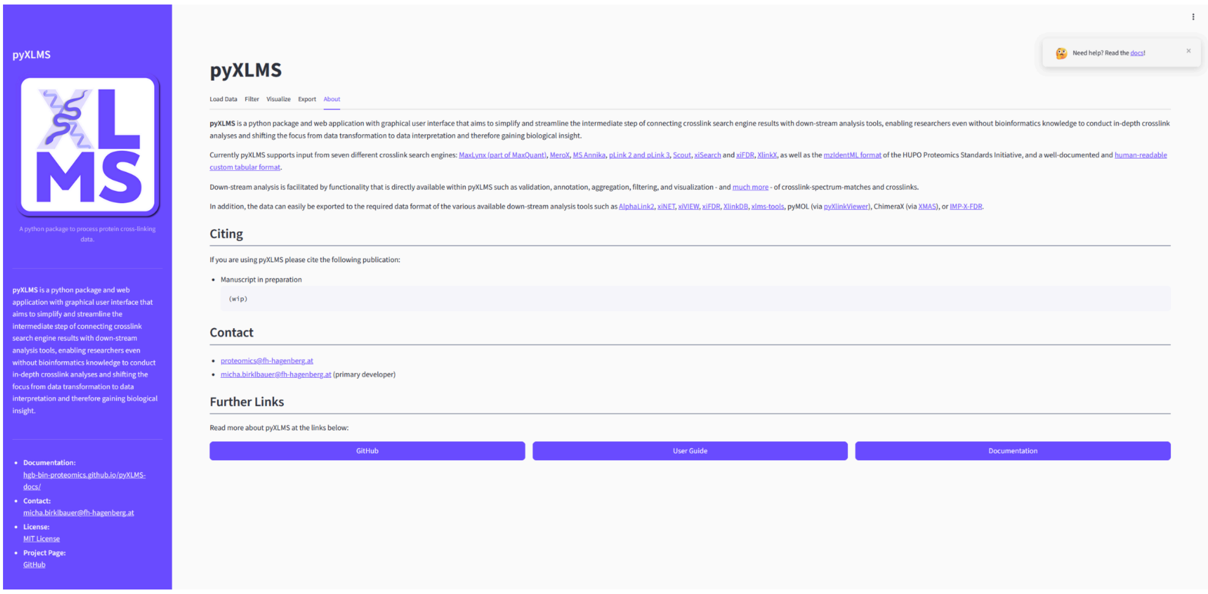


TIP

*If you run into any problems with the pyXLMS web application or have any questions, please check our [help page](#) on how to get support!*
